# Statistical properties of simple random-effects models for genetic heritability

**DOI:** 10.1101/087304

**Authors:** David Steinsaltz, Andrew Dahl, Kenneth W. Wachter

## Abstract

Random-effects models are a popular tool for analysing total narrow-sense heritability for simple quantitative phenotypes on the basis of large-scale SNP data. Recently, there have been disputes over the validity of conclusions that may be drawn from such analysis. We derive some of the fundamental statistical properties of heritability estimates arising from these models, showing that the bias will generally be small. We show that that the score function may be manipulated into a form that facilitates intelligible interpretations of the results. We use this score function to explore the behavior of the model when certain key assumptions of the model are not satisfied — shared environment, measurement error, and genetic effects that are confined to a small subset of sites — as well as to elucidate the meaning of negative heritability estimates that may arise.

The variance and bias depend crucially on the variance of certain functionals of the singular values of the genotype matrix. A useful baseline is the singular value distribution associated with genotypes that are completely independent — that is, with no linkage and no relatedness — for a given number of individuals and sites. We calculate the corresponding variance and bias for this setting.

**MSC 2010 subject classifications:** Primary 92D10; secondary 62P10; 62F10; 60B20.

## 1. Introduction

Genome-Wide Complex-Trait Analysis, known as GCTA, introduced by Jian Yang and collaborators in 2010 in [53] has led both to a profusion of research findings across the biomedical and social sciences and to exuberant controversy. Use of the method has leapt ahead of clarity about its statistical properties. There has been extensive discussion of sensitivity to violation of assumptions, but no consensus on performance when its most basic assumptions are satisfied. Is statistical bias a concern even in the simplest settings? Some say yes [20]. Some say no [55]. Do tractable standard errors depend on the presence of residual population stratification? Formulas in the literature [46] leave the answer murky.

In this paper we seek to settle these basic questions.

- For “Simple GCTA”, defined below, with *n* respondents and *p* genetic markers, we go beyond order 
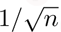
, derive fromulas for bias to order 1/*n*, and show this intrinsic bias to be negligible in practice.
- For less simple settings, we analyze two known sources of bias:

– negative estimated values for the GCTA parameter representing heritability, which we argue remain meaningful within the model and should not be excluded;
– causal markers at fixed rather than randomly distributed subsets of sites, introducing potential bias which we characterize and bound.
- We present interpretable expansions of standard error drawing on eigenvalue theory, depicting the contrasts between standard errors in the absence and in the presence of population stratification.

Our results are directed at cases in which model assumptions hold true. Effects of model misspecification, widely treated in the literature, are considered briefly to round out our study.

The data for GCTA are assays of very large numbers of Single Nucleotide Polymorphisms (SNPs) in the genomes of individuals along with measurements of putatively heritable traits. The model of GCTA is an example of a “linear mixed model” in which the contribution of SNPs to trait values are treated as random effects. (We do not consider fixed effects in this paper.) Mixed models form a natural framework for the estimation of total heritability for traits whose variability is determined by a wide variety of sites, rather than by specific identifiable SNPs each with strong influence. GCTA found notable application in work by Yang and Visscher and associates to identify “missing heritability” in height [53, 50] and other complex traits, and many groups have followed their lead. The popular algorithm for implementing GCTA goes under the acronym GREML, for “Genome-Wide Restricted Maximum Likelihood”. We limit our attention to the random-effects portion of the model — the same arguments would apply to the reduced phenotypes after we eliminate fixed-effects — so there is no “restriction” to make, and we are considering simple maximum likelihood estimates; we employ the general term GCTA here rather than GREML. An alternative estimation method for the same model, under the name “LDSCORE Regression”, is also coming into wide use, bringing up as yet unexplored statistical issues paralleling those we examine here for GCTA.

The goal of these methods is to estimate total additive heritability, without the overfitting that arises in attempts to identify specific loci influencing a trait. As we have said, most examination of pitfalls – including two papers [51, 58] with “pitfalls” in the title – have emphasized issues that arise from kinds of model misspecification: from single large-effect alleles (better treated as fixed effects), from reliance on linkage disequilibrium when causal alleles themselves are unobserved [42, 22, 43], from nonlinear increase in heritability estimates with increasing numbers of SNPs (reflecting saturation of coverage of a smaller subset of genuinely causal loci), and from ascertainment bias in binary traits [24, 4, 12].

Recently, Kumar et al. [20] have taken a different tack, criticising GCTA on statistical grounds, on what might be considered issues of inherent mathematical fallibility for estimation in models relying on high-dimensional covariates. Their paper has elicited rebuttals from Yang et al. [55] and [56] and rejoinders to the rebuttal [21] and [19] from Kumar et al. Parts of that exchange are devoted to GCTA in more complicated settings, but our results in this paper settle some of the points in contention.

In “Simple GCTA”, as we use the words, all causal SNPs are observed, all observed SNPs are causal, and the sizes of causal effects are all drawn independently from the same centered normal distribution. We abstract away from the need, important in practice, for tagging unobserved causal alleles by observed alleles taking advantage of linkage disequilibrium. Non-genetic variance is contributed by independent draws for each sample from another centered normal distribution. Fundamental statistical properties are brought into the spotlight by studying this streamlined version.

For us, the matrix of genotypes is fixed and observed without error; we condition on it. Statistical properties of GCTA estimates depend on the matrix through its squared singular values, eigenvalues of the Genetic Relatedness Matrix. Numbers of samples *n* are assumed to be in the tens or hundreds of thousands, and numbers of SNPs *p* much larger; the ratio *μ* of samples to SNPs is a key parameter.

Empirical studies have alluded to or reported a wide variety of patterns for sets of squared singular values, sometimes concentrated near unity, sometimes dispersed across orders of magnitude. We consider two extremes of contrasting settings intended to bracket the reported patterns in the literature. First, (A), is the “independent setting”, with a genotype matrix resembling one random draw from an ensemble of matrices with independent entries, thereby assuming an absence of population stratification and of linkage disequilibrium. Second, (B), is a “stratified setting”, represented in this paper by three flavors of stylized distributions, evoking genotype matrices whose singular values suggest deep population stratification.

While the independent setting is a frankly artificial idealization, it serves as an indispensable guide. Intuition suggests that the information content from a set of SNPs in linkage disequilibrium should resemble the information content from a smaller set of independent SNPs. The independent setting with a downward-adjusted ratio of SNPs to samples gives a good starting point for realistic genotype matrices. With this advantage in mind, we devote special attention to explicit formulas for estimator bias and variance in the independent setting

The key tool in our statistical analysis of GCTA is a profile likelihood function that reduces the estimation problem to finding the root of a univariate function as well as simplifying simulations. The model is set out in Section 2 and the profile likelihood and bias formula are derived in Section 3 under the assumptions of Simple GCTA. Sections 4 and 5 take up the more complicated issues associated respectively with negative estimates of the heritability parameter and with subsets of causal SNPs and other aspects of model specification. Section 6 derives the formulas relevant to the independent setting, and Section 7 sums up.

It is common to truncate estimates of heritability at 0. This produces a bias, which can be estimated from the normal approximation. There is a question, though, whether this truncation is appropriate. Retaining and interpreting negative estimates is an attractive alternative. Section 4 puts forward the claim that GCTA naturally resides within a statistical model in which the parameter playing the role of heritability can be negative and still meaningful. Although heritability as ordinarily defined, being the ratio of two variances, cannot be negative, the parameter in the statistical model admits another interpretation, one with precedents in genetics going back to the 1940s and before. We develop the implications of this point of view and outline a biological story of how negative values can legitimately arise.

With regard to subsets of causal SNPs, Section 5, under the heading of “Model Misspecification”, assesses potential biases at several levels of complexity. Yang et al. remarks at the outset of [53] that it is harmless to relax the assumptions of Simple GCTA to allow non-zero effect sizes to be confined to an unknown random subset of observed SNPs. We agree with this claim up to a point, but show that it needs some correction. As Yang et al. recognize, non-zero effects solely on a **fixed**, unidentified subset of SNPs can in principle, as we show, introduce a bias of arbitrary size and direction. While it is true (as we confirm, using more direct arguments) that this bias averages out close to zero when the causal set is considered as a randomly chosen subset of all SNPs, this bias increases the expected error in the heritability estimates. Under strong but reasonable additional assumptions, however, we can estimate this additional error, and show that it is small under most circumstances. Non-zero effects on unobserved causal SNPs in linkage disequilibrium with observed SNPs is a complicated, widely-discussed issue, on which we offer some brief views of our own in Section 5.4.2.

Our findings about bias and variance run contrary to conclusions of Kumar *et al*. in [20]. In particular, we see the limitations on accurate GCTA in the absence of stratification arising not from bias but from large standard errors, in contrast to their conclusions about the salience of bias. We see population stratification in small doses enhancing the accuracy of GCTA estimates of heritability — at a cost in interpretability — in contrast to their general contention that stratification undermines the stability of estimates. However, we do see large departures from the independent setting imperilling the accuracy of GCTA, leading us to be less sanguine than Yang et al. [55] about the statistical properties of these genome-wide random effects models.

Many further issues about GCTA arise in more complicated settings under more flexible assumptions and rightly engender continuing debate. However, given the impact of increasingly complex generalizations of simple GCTA to test association [18, 17, 61, 27, 40, 45, 37, 31, 30, 29, 26, 62, 3], partition heritability [59, 54, 9, 2], predict phenotype values [39, 60, 41, 6], and learn “co-heritabilities” [44, 25, 5, 7], it is important, if possible, to reach consensus on some core facts. Our analysis of bias and variance for simple GCTA aims at this goal.

## 2. The GCTA model

We suppose we are given a data set consisting of an *n* × *p* matrix *Z*, considered to represent the genotypes of *n* individuals, measured at *p* different loci. There is a vector **y**, representing a scalar observation for each of the *n* individuals. The underlying observations are counts of alleles taking the values 0, 1, 2, but the genotype matrix is centered to have mean zero in each column and normalized to have mean square over the whole matrix equal to 1.

It is common practice to go further and normalize each column to have unit variance either empirically or under Hardy-Weinberg equilibrium. (SNPs far from Hardy-Weinberg equilibrium are generally excluded by quality control procedures, so these two alternatives amount to much the same thing.) Such normalization, usual in GCTA, gives all columns equal weight in producing genetic effects. This assumption that normalized SNPs have i.i.d. effect sizes implies that unnormalized SNP effect sizes decrease with increasing allele frequency [13, 48]. Except where noted, we do not assume this column-by-column normalization. The normalization of the sum of squares of the whole matrix is required to make sense of the interpretation of the parameter representing heritability.

The basic assumption of the model is the existence of a random vector **u** ∈ ℝ^*p*^ of genetic influences from the individual SNPs such that

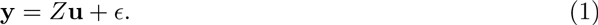

The vectors **u** and *ϵ* are assumed to be independent and to have zero means and i.i.d. normal components. The variances are determined by two parameters, which are to be estimated: *θ* represents the precision (reciprocal variance) of the non-genetic noise and *ψ* represents the heritability, entering the model as the ratio of genetic variance to total variance. We use *ψ* rather than the more conventional *h*^2^ in order to accommodate possibly negative estimates, which figure in Section 4.3. It will be convenient at some points to use the parameter *ϕ* = *ψ*(1 − *ψ*) in place of *ψ* itself.

Writing (*θ*_0_, *ψ*_0_) or (*θ*_0_, *ϕ*_0_) for the true values from which the data are generated, we have 
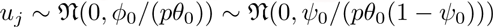
 and 
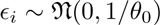
.

In the discussion by [20], much weight is placed on *Z* being a “random” matrix. There are several senses in which *Z* may reasonably be thought of as random:

1. The individuals are sampled from a larger population.
2. The SNPs have been selected from a larger set of possible SNPs.
3. The genotypes of individuals have been formed by random processes of mating, mutation, selection, and recombination.
4. There are random errors in the genotypes.

None of these substantially affects the analysis we carry through in this paper (although, with regard to the fourth kind, see Section 5.3). The model assumes that all genetic causality runs through *Z*, so that for purposes of estimation it may simply be taken as a deterministic known quantity, as is standard for covariates in regression models. On the other hand, as in standard linear regression models, some choices of independent covariates make the regression problem easier than others, so it is worth considering which *Z* may be “likely” to occur.

As we show shortly — and as [20] correctly point out — the properties of this statistical method are determined entirely by the singular-value spectrum of *Z*. In our independent setting (A), for a population without stratification and linkage disequlibrium, the empirical distribution of the singular values is expected to be close to a known limiting form, featured in Section 6 and depending only on the dimension ratio *μ* = *n*/*p*. In our stratified setting (B), with a proportion of singular values orders of magnitude larger than those typical of setting (A), qualitative generalizations need not rely on the detailed singular value spectrum.

## 3. The profile likelihood and bias formula

### 3.1. The likelihood function

We begin our derivation of expressions for bias and standard error in estimated heritability by transforming GCTA likelihood estimation from a two-parameter to a one-parameter problem. We define a profile likelihood function whose maximization only requires finding the zero of a univariate function. This device is the key to the derivation, and it offers the bonus of facilitating simulations. By standard likelihood arguments, to order 
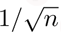
 estimator bias is zero. But 
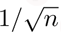
 is not small enough, in many GCTA applications, to make the next term, of order 1/*n*, negligible, if it comes with a large constant. We need to compute the next term in order to resolve the conflicting claims about bias in the literature.

The expression for bias brings with it an expression for estimator variance to order 1/*n*, equivalent to the standard but unwieldy expression from Fisher Information, given, e.g., on pages 234–235 of [46], We go on to make bias and variance interpretable by expanding them in the independent setting in terms of dimension ratios and comparing with stylized cases for the stratified setting.

Conditioned on **u** and *Z*, and defining the Genetic Relatedness Matrix or GRM *A*:= *p*^−1^*ZZ*^*^, the measurements **y** are normally distributed with mean zero and covariance matrix

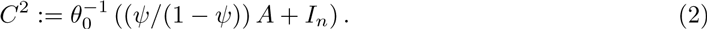

Let *Z* = *U*diag(*s_i_*)*V*^*^ be the singular-value decomposition of 
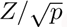
, and rotate the observations to diagonalize the covariance matrix, obtaining

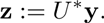

The elements of **z** are independent centered normal random variables with variances 
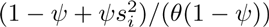

The log likelihood is then

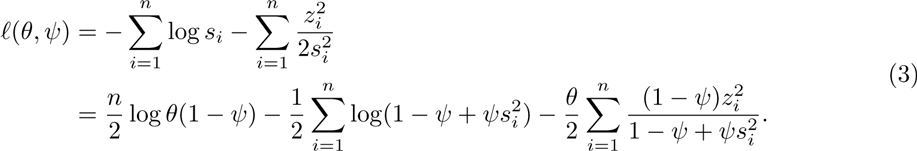

Note that **z** depends only on *Z* and **y**, not on the parameters.

We observe here that [20] claim (without demonstration) that the presence of singular values in the denominator of the likelihood creates “instability” in estimates based on this likelihood when the singular values are small, and that the dependence on the projection onto left singular vectors creates instability when the singular values are close together. Neither is true. Their representation of the log likelihood differs from the one we have here by the addition of the log determinant of *A*. This is a very large number if there are singular values close to zero (and indeed infinite if singular values are zero), but the addition of a constant, however large, has no influence on likelihood-based estimation. (We note here that the work done by Sylvester’s Theorem (their equation [A6]) is unnecessary as soon as we interpret the determinant as a product of squared singular values. Furthermore, in the case when a singular value is exactly zero, which occurs automatically when using de-meaned SNPs, the conditions for applying Sylvester’s Theorem are not satisfied.) Similarly, under the assumptions of the model, the *z_i_* are independent normal random variables, with variances 
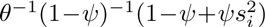
, which is positive so long as the environmental contribution *θ*^−1^ is positive.

### 3.2. Solving for the MLE

The partial derivatives of the log likelihood are

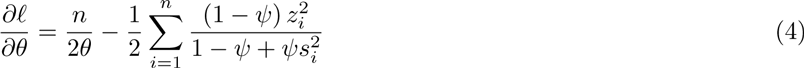

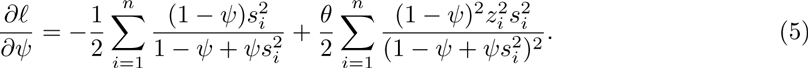

Solving *∂ℓ*/*∂θ* = 0, we get

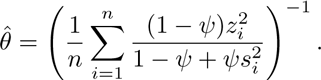

Substituting into (5) we get the profile likelihood

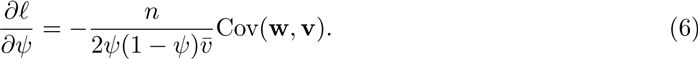

Here

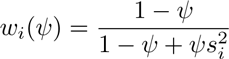

and

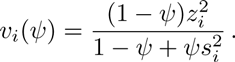

The *w_i_* are not random, whereas for each value of *ψ* the 
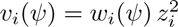
 are random variables. At *ψ* = *ψ*_0_, the expected values of *θ*_0_*v_i_* for all *i* are unity.

Note that the normalization that makes all columns of *Z* sum to 0 induces a singular value of 0, which we might as well place at the end of the list as *s_n_* = 0. This corresponds to the constant left singular vector 
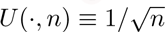

The symbol Cov in equation (6) represents the empirical covariance of the elements of the n-dimensional vectors **w** and **v**. Similarly, we use Var with a vector argument for the empirical variance of the elements. When the vector elements are themselves random variables, as in this case, the output of Cov and of Var are themselves univariate random variables.

We define *τ_k_* to be the empirical *k*-th central moment of the elements of the vector *w_i_*(*ψ*_0_); that is,

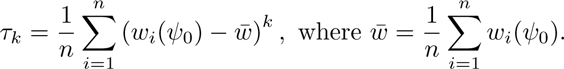

(Note that *τ*_1_ = 0, and we set *τ*_0_ = 1.) Since *w_i_* ∈ [0, 1], we know 
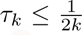
, and also 
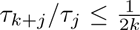
; for *k* ≥ 3 we may replace 2*k* by 2*k* + 2 (*cf*. [8]). Formulas for *w̄* and *τ*_2_ and *τ*_3_ under assumptions about the singular values of the genotype matrix *Z* are derived in Section 6. Bear in mind that, in the current model, **w** is a function of the genotypes, so it is not random. The only randomness is in **u** and **y**, hence in **z**, hence in **v**.

We have now arrived at our profile likelihood equation. Setting the left-hand side of equation (6) to zero, we have the maximum likelihood estimate of *ψ* equal to a root of the univariate equation

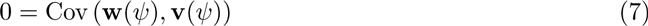

The estimate is a function of the transformed observations **z** and of the singular values *s_i_* of the scaled genotype matrix 
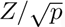
.

Our next task is to express equation (7) in terms of a power series in the differences *ψ* − *ψ*_0_. The right-hand side of equation (7) as a function of *ψ* for any fixed realization of the random quantities 
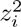
 is a polynomial in *ψ* times a weighted sum of fractions 
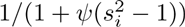
 and it is an analytic function everywhere except on that part of the real axis outside the interval 
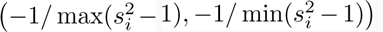
, which includes the interval [0, 1] since the mean of the squared singular values is unity. We assume now that *ψ*_0_ ∈ (0, 1) and define

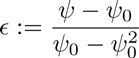

(Implications of allowing *ψ* to be negative will be discussed in section 4.3.) Then the right-hand side of equation (7) has a convergent Taylor series expansion for *ϵ* in an interval around the origin.

We begin with the identities

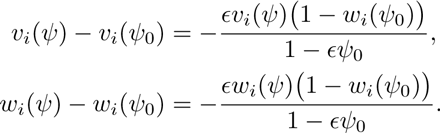

Rearranging terms yields

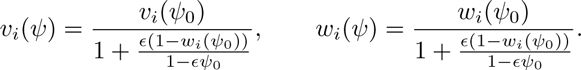

We now work out explicit formulas for coefficients in the Taylor expansion of the score equation up to second order in *ϵ*. (This expansion is the usual one for expressing asymptotic variance in terms of Fisher information, but we need explicit formulas for the sake of our explicit approximation for the bias and for a similar expansion in section 5.4.) By the bilinearity of covariance,

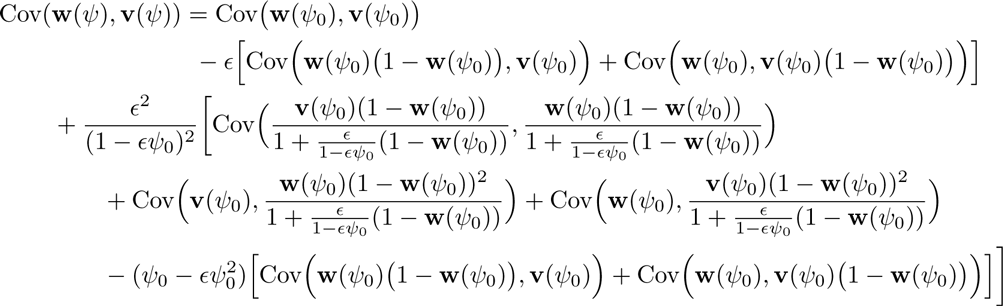

At the maximum likelihood estimate *ψ* = *ψ̂*, the left-hand side is zero. So, for the value of *ϵ* corresponding to *ψ̂*, we obtain

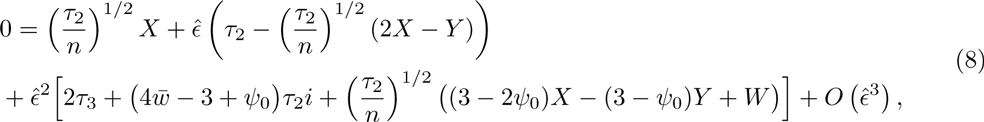

where

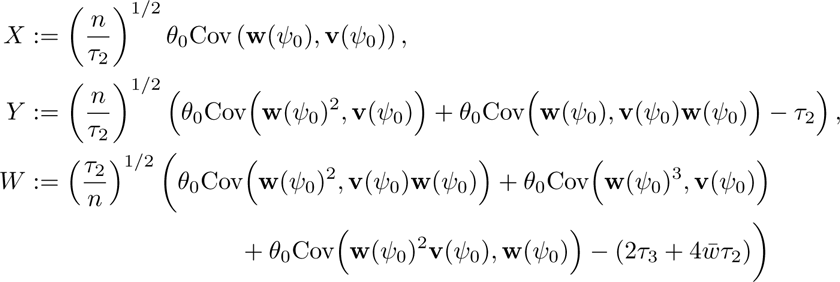

We observe now that *θ*_0_**v**(*ψ*_0_) is a vector of i.i.d. 
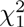
 random variables. It follows that *X*, *Y*, and *W* (which are close to normal random variables) have expectation values of zero and product moments

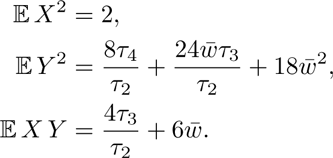

(Corresponding formulas for *W* are of the same sort, but longer.)

We now examine the powers of 
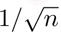
 that enter into our equation for *ϵ̂*. For very large *n*, *ϵ̂* is of a magnitude comparable to −*νX*, where

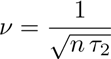

We rewrite equation (8), after dividing by *τ*_2_, in the form

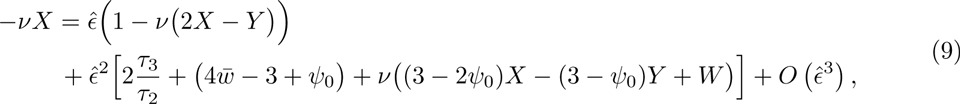

By Reversion of Series, expanding in powers of *ν*, we have

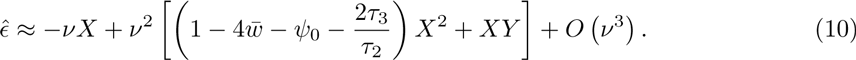

For large *n* and small *ν*, this expansion is valid when the values of *X*, *Y*, etc. are not so extreme as to prevent the score equation from having a unique root. The probability of extreme values can be made small by taking *n* to be large. It suffices to take 
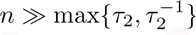
, so that *ν* ≪ 1, along with 
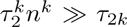
 for positive integers *k*, but the formulas for moments of *ϵ̂* remain accurate for many cases with small *τ*_2_ and *ν* > 1.

### 3.3. The bias formula

Our formula for bias to order 1/*n* follows by taking expectations of both sides of equation (10).

- Confirming standard results, we see that to first order in *ν* (i.e. to order 1/*n*^1/2^), the estimator *ψ̂* is unbiased.
- There is non-zero bias on the order of *ν*^2^, i.e. 1/*n*. If we drop terms of order *ν*^3^ and higher, we get

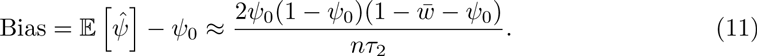
- Squaring both sides of equation (10), we confirm the formula for estimator variance,

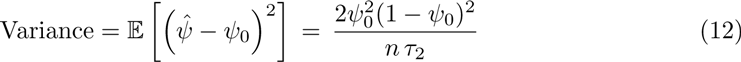

The mean of the eigenvalues 
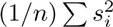
 is the trace of the matrix (*pn*)^−1^*ZZ*^*^, which is 1/*n* times the sum of the squares of the entries of *p*^−1/2^*Z*. As described in section 2, we are assuming that the genotype matrix has been normalized to make this mean equal to 1. Jensen’s Inequality then implies

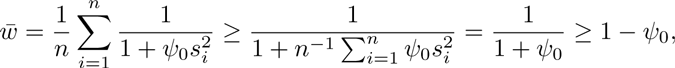

To order 1/n, the bias is negative.

The ratio of bias to standard error for *ψ̂* is

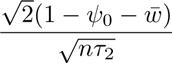

The ratio will be small when *nτ*_2_ is large.

When the variance in 
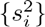
 is small we have delta-method approximations

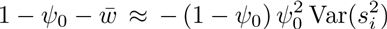

and

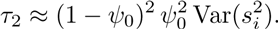

The consequence is an approximate bias in *ψ̂* of (−2*ψ*_0_/*n*). These approximations may fail utterly if eigenvalue variance is large, as it may be in stratified settings, but they correctly pick out leading terms when eigenvalue variance is small.

For the the independent setting, we can be more precise. Exact formulas for the contributions of order 1/*n* to bias and variance in estimated heritability and for the moments of *w_i_* for the independent setting are derived in Section 6, along with expansions up to second order in *μ* = *n*/*p*. The expansions give

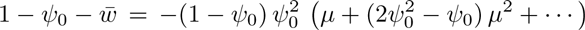

and

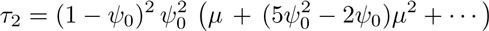

Estimator bias is given by

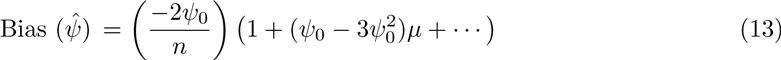

Estimator variance is given by

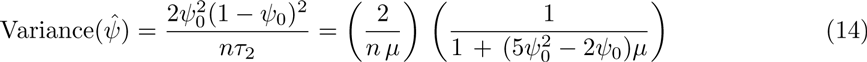

We conclude that in the independent setting bias is indeed a very small negative number, a third-decimal effect even for samples no bigger than a thousand respondents. Variance, on the other hand, increases as the number of SNPs per person 1/*μ* increases, implying standard errors as large as 0.14 with 10, 000 people and a million SNPs.

The contrasting “stratified setting”, as we are using the term, embraces a wide range of alternatives found in empirical cases whose genotype matrices have a subset of singular values substantially larger than singular values from the independent setting. Since the mean of squared singular values is constrained to be unity, each large singular value must be balanced by a number of small ones. We review the behavior of bias and variance under three stylized models incorporating such balance and broadly resembling singular value distributions described in the literature.

The first two models are built from a specification with paired point masses developed in Section 6, namely a distribution for 
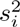
 which puts mass *β*/(*β* + 1) at 1/*β* and puts mass 1/(*β* + 1) at *β*.

In the first stylized model, a moderately small proportion *α* of squared singular values are drawn from this paired-mass distribution, while the remaining 1 − *α* recapitulate those from the independent setting. Means and mean squares for *w_i_* are weighted averages of expressions given in Section 6. Such a “dosage” of more widely dispersed singular values does increase *τ*_2_ and reduce the standard error of estimation, but not by much. Small *α* even in combination with large *β* limits the improvement. When *ψ* = 1/4 and *α* = 1/100 with **μ** = 1/25, standard errors drop by less than 25%. The dosage also shifts *w̄*, but the bias remains very small in comparison to the standard error for typical, sizable *n*.

In the second stylized model, the cluster of singular values close to unity characteristic of the independent setting is taken out, putting *α* = 1 and leaving the paired-mass distribution on its own. For large *β*, the variance of the squared singular values is close to *β* and the moments of *w_i_* given in section 6 lead to approximations for estimator bias and variance:

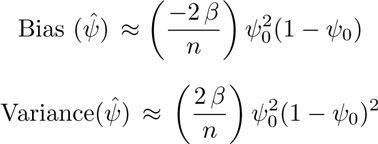

Here, the larger the variance in squared singular values (roughly *β*), the larger the variance in *ψ̂*. Instead of affecting estimator accuracy in the same fashion as *μ* = *n*/*p* (the variance of 
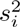
 in the independent setting) *β* takes on a role like 1/*μ*, eroding accuracy as it grows. The stylized setup makes the reason apparent. Large squared singular values have to be balanced by large numbers of near-zero values to preserve the mean of unity. The transformation from *s_i_* to *w_i_* discounts the leverage of large values while the preponderance of small ones pull *w̄* toward unity and *τ*_2_ toward zero.

The third stylized model posits squared singular values from a lognormal distribution, mimicking the appearance of singular value graphs in [20] and elsewhere. Denoting the variance of the squared singular values by *γ*, under the constraint of unit mean the variance of the underlying normal is log(1 + *γ*) and its mean is −(1/2) log(1 + *γ*). Our *w_i_* then follows a so-called “logit-normal” distribution. Closed-form moments for *w_i_* are not available, but their behavior is easy to infer. As *γ* increases, squared singular values become heavily concentrated near zero and *w_i_* near 1 − *ψ*. The variance of *w_i_*, that is, *τ*_2_, increases to a maximum and then falls off slowly toward zero. Moderate stratification reduces standard errors of estimation, but large departures from the independent setting raise standard errors and spoil estimates.

### 3.4. Implications of the formulas

The formulas of Section 3.3 have implications that may at first seem surprising but have logical explanations. First, in the context of Simple GCTA, when we are dealing with independent random genotypes, increasing *p* — providing more data — seems to make accurate estimation harder. For fixed *n*, standard errors go up with *p*, not down. Second, stratified populations — something that would ideally be avoided in real data [53], and that the earlier analysis of [20] suggested would undermine the model even in theoretical, unconfounded data— seem up to a point to alleviate the problem of large standard error.

In fact, neither of these implications is surprising. The apparent paradox of more data producing a worse estimate dissolves when we recognise the structure of GCTA. Increasing *p* does not simply provide more data. It changes the assumed set of influences on the phenotype. In Simple GCTA, increasing *p* means dividing the same overall genetic effect into tinier pieces. In more complicated versions, where causal SNPs comprise a subset of (observed or unobserved) SNPs, as discussed in Section 5.4, positing larger numbers of causal SNPs similarly means dividing up the overall effect. The Law of Large Numbers tends to equalize genetic endowments among individuals, regardless of which particular SNPs happen to have the largest effects. Naturally, the situation becomes more complex when causal SNPs are sparse and linkage disequilibrium is crucial, issues that figure prominently in the literature.

It is also no surprise that some degree of population stratification, as it augments the variance of squared singular values, reduces standard errors of estimation. Zero or near-zero singular values reflect sets of individuals with high genetic similarity. Their presence allows noise to be most easily isolated from the genetic effect. It is common practice to study twins to simplify the identification of genetic effects. The reason behind the standard practice of removing relatives from a sample and pulling out population strata as fixed effects is a belief that relatives are likely to have their common genetic influences confounded with shared environment and that genetic strata are likely to reflect stratification in non-genetic respects, hence also create confounding [38, 36, 53]. The reason is not that such individuals make the statistical analysis inherently difficult.

The GCTA model implies a correspondence between the covariances of the high-dimensional genotype and the covariances of the measured trait. Clearly, there is information in *Z* and **y**, but it is not immediately apparent where it is and how much is there. Passing to the diagonalization of *Z*Z* clarifies the situation: The information is in the different magnitudes of the rotated components. This is very diffuse information. We are faced with a large number of observations *z_i_* from very similar distributions, differing only in their variances, which are 
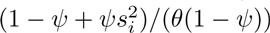
. We are trying to identify *ψ* as the parameter that best orders the *z_i_* by magnitude. It is apparent that the challenge increases as the *s_i_* become more compressed, as our formulas provide.

### 3.5. A symmetry relation

The demonstration in section 3.2 that the GCTA estimator of heritability is approximately unbiased depends on an approximation to order 1*/n* that is incomplete from a practical point of view, insofar as we do not show that terms of higher order than 1/*n* are genuinely smaller for ranges of parameters of interest.

The demonstration can be strengthened by appeal to a symmetry relation. If we replace **y** by **ỹ** = *A*^−1/2^***y***, then we obtain a sample from the same class of multivariate normal distributions defined in (2), with the matrix *A* replaced by *A*^−1^, and the parameter *ψ* is replaced by 1 − *ψ*. The parameter *θ* becomes *θ*(1/*ψ* − 1).

A genotype matrix that would produce *A*^−1^ would not have the usual properties of the normalized genotype matrices we have been considering. Any claim that the GCTA model generally yields estimates biased in one direction must depend on such properties. In principle an allowable data set that yields a positively biased heritability estimate can be paired with an alternative allowable data set (obtained by inverting the GRM) that yields a negatively biased heritability estimate, without changing the form of the model.

## 4. Interpreting the heritability parameter

As Albert Jacquard [15] pointed out decades ago, *narrow-sense heritability* —— *h*^2^, the parameter that we have designated here for notational convenience as *ψ* —— has conventionally two distinct meanings:

1. The proportion of total variance attributable to additive genetic effects;
2. The slope of the linear regression line of a child’s phenotype measurement on the mean of the parents’ measurements.

Both meanings appear in the earliest works to give a quantitative operational definition to *heritability*, in particular [32]. (For more about the history of the notion of heritability see [1].)

The correspondence between these two meanings depends on the additive model, where genetic and non-genetic effects are independent and sum together to produce the phenotype. When we have general genetic relatedness (rather than parental relations with fixed 50% relatedness) heritability is analogous to a regression coefficient relating phenotypic similarity to genotypic similarity. We describe in section 4.1 how this analogy is made precise. We go on in section 4.3 to explore the relevance of these observations for the problem of negative estimates of the heritability parameter, a statistical embarrassment if heritability is to be interpreted as a ratio of variances, but a natural outcome under a regression-based definition.

### 4.1. GCTA as a linear regression model

The GCTA model has been formulated as a random-effects model, but it is equivalent to a multivariate normal model written out in section 2. In this section we describe how the model may also be understood as a linear regression model. In their original paper [53], Yang and coauthors spell out an analogy between GCTA and a different form of linear regression. They regress squared trait differences between pairs of individuals on corresponding elements of the Genetic Relatedness Matrix, with *n*(*n* + 1)/2 points and correlated errors. Instead, we draw an approximate comparison between GCTA and regression with *n* points and independent errors.

Under the GCTA model, we have observations

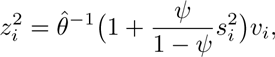

where *v_i_* are i.i.d. chi-squared random variables with one degree of freedom. When 
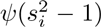
 are uniformly small, we may write this model as

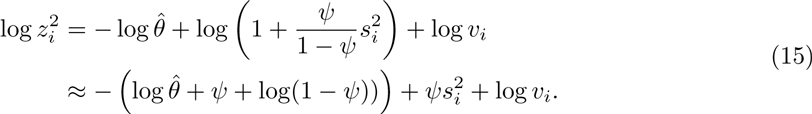

If we compare this to a standard linear regression problem (with log 
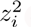
 as the dependent variable, 
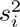
 as the independent variable, and log *v_i_* as the noise), we would expect to have variance of the slope estimate inversely proportional to the variance of the independent variables. That is, when the 
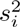
 are tightly clustered, there will be large errors in the estimate of the slope, which is the parameter that gives information about *ψ*.

In addition, we would have to cope with the fact that the noise term is not normal, but log chi-square.This is highly left-skewed, with a very short tail on the positive side, and a long tail (asymptotically exponential) on the negative side.

### 4.2. Simulations

As an example, we plot in figure 1 an example based on the same dimensions as those for the genotype matrix in the celebrated paper [53], but drawing singular values from the independent setting, not from the (unreported) empirical distribution of singular values underlying that study. For *n* = 4000 and *p* = 100000, we show a scatterplot of the pairs 
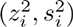
 that would be obtained from the singular-value decomposition of a genotype matrix with the appropriate dimensions. Note that the lines are very similar, and have relatively little leverage relative to the huge scatter in the values of 
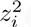
.

**Fig 1.**
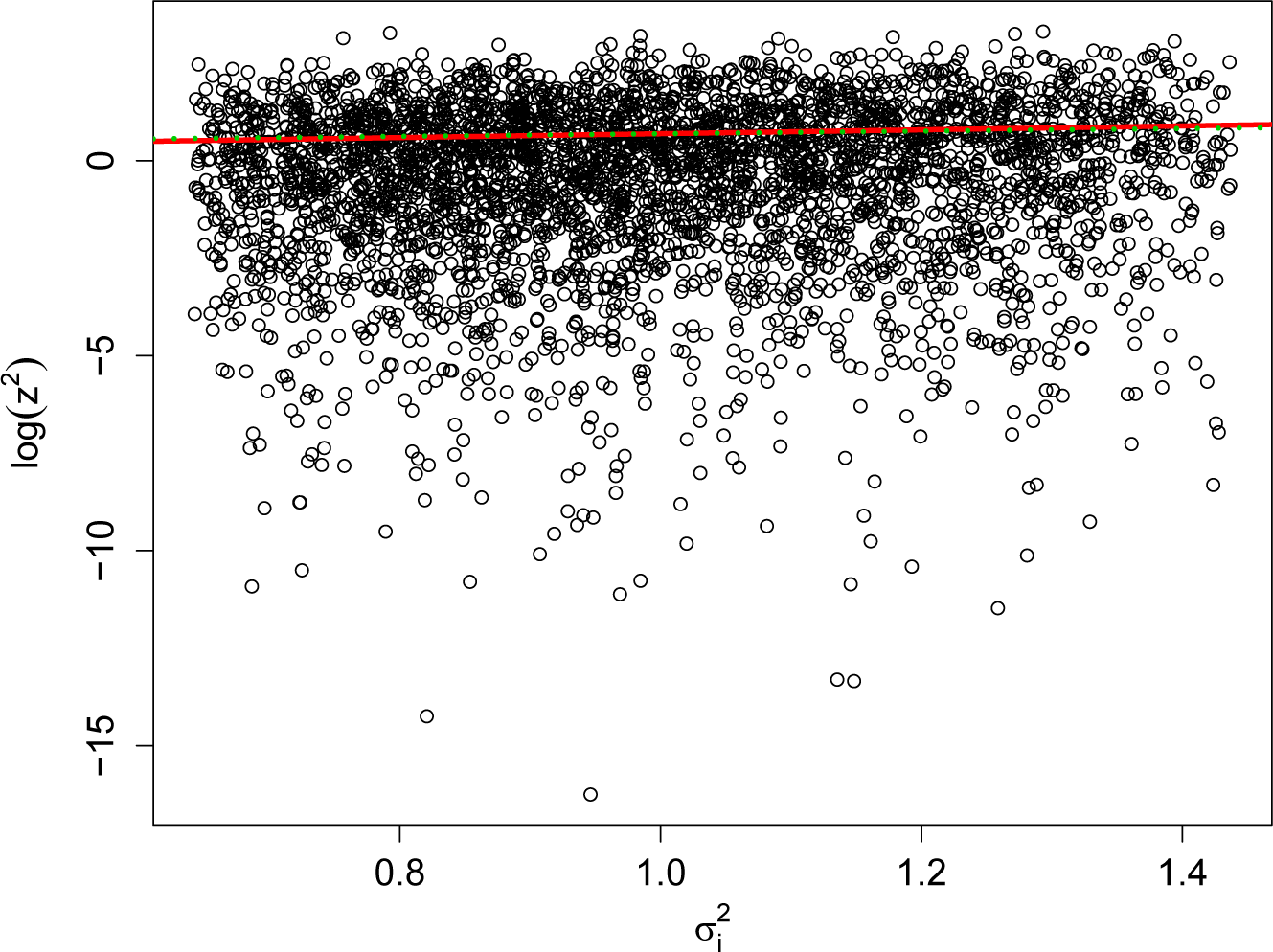
Linear-regression approximation of the estimation problem for *ψ*. The solid red line shows the correct line, corresponding to *ψ* = 0.5, while the dashed green line corresponds to *ψ* = 0.25.

**Fig 2.**
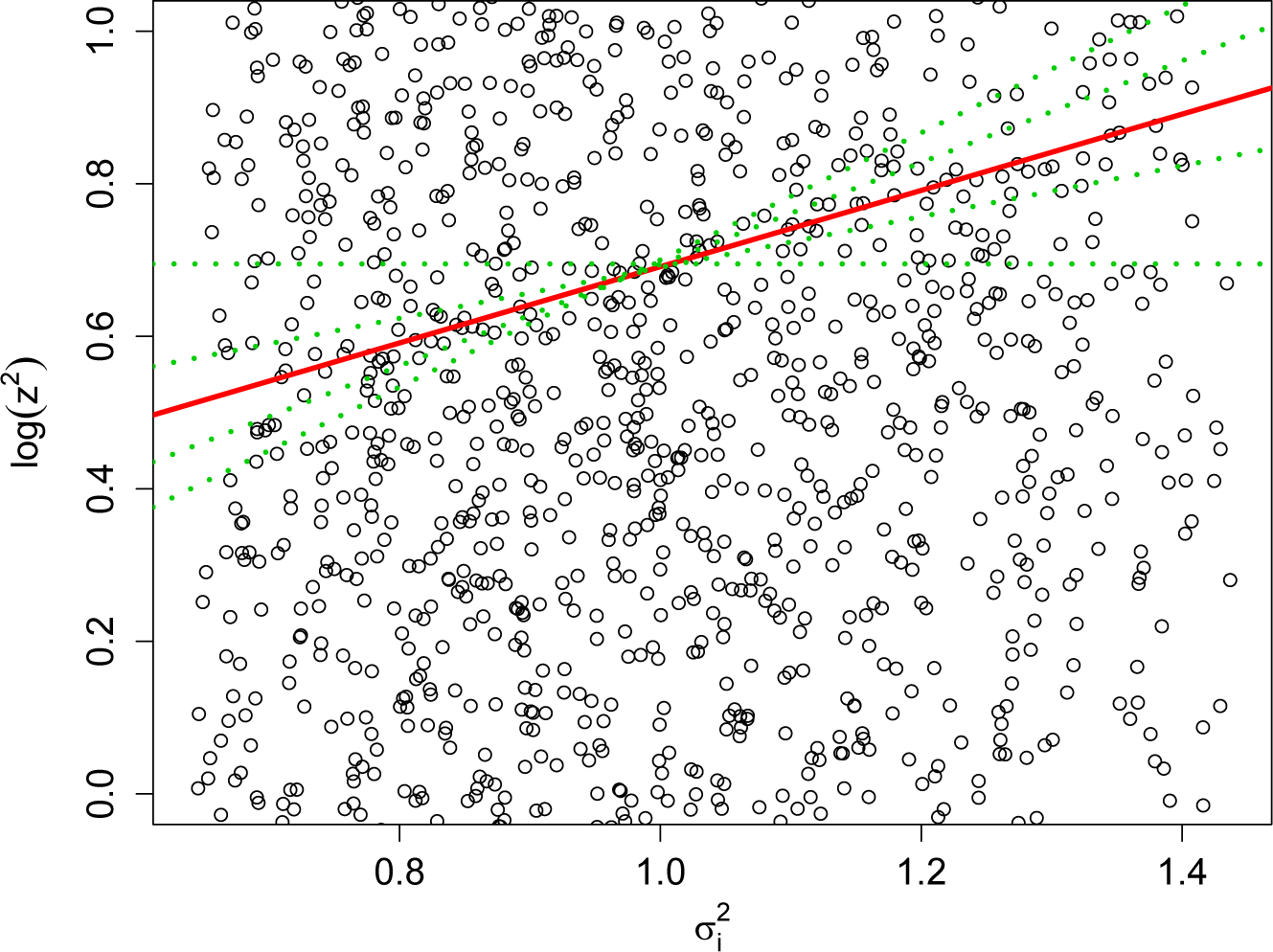
Blow-up of the linear-regression approximation, showing just the range 
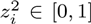
. The dashed green lines correspond to *ψ* = 0, 0.33, 0.67, 0.84.

In the independent setting, the known limiting measure for the singular values presented in Section 6 simplifies the task of exploring the GCTA model through simulations. Instead of needing to perform matrix multiplications and diagonalizations on a random *n* × *p* matrix, where *n* and *p* may be on the order of 10^5^ or 10^6^, which quickly becomes computationally expensive, we may instead start with the singular values. These are *n* equally spaced values from the singular value distribution, to which we contribute *n* normally distributed random variables *z_i_* with mean zero and variance 
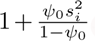
. We solve for *ψ̂* that makes the vector (*z_i_*) and the vector (*w_i_*) uncorrelated. That is, Cov(*z_i_*(*ψ*), *w_i_*(*ψ*)) is a univariate function of *ψ* that is easy to compute, and will typically cross zero exactly once. (The exceptions are when the covariance is strictly negative for all *ψ*, meaning that the likelihood is increasing, so that the MLE is *ψ̂* = 1; and when the are multiple solutions.) The simulations above, and in section 4.3, have been carried out by this procedure. This should be essentially identical to the results that we would obtain from simulating a random GRM and a random trait vector. (Furthermore, we could simulate a different model for the GRM simply by choosing a different distribution for the singular values.)

### 4.3. Negative heritability

The parameter *ψ*, corresponding to narrow-sense heritability, may be estimated as negative. Negative estimates of the heritability parameter are often dismissed as a mathematical abstraction, values in a range that arises purely formally and that may only be reported for formal purposes, as part of an ensemble of estimates that collectively are unbiased. Several recent studies [57, 2, 9, 63] have reported individual negative heritability estimates in this way, including them in averages that themselves came out positive. In [16] a point estimate of –0.109 is obtained for heritability of horn length in Soay sheep. It is immediately dismissed with the statement that “it is impossible to have negative heritability” and the inference is drawn that the true heritability must actually be a small positive number in the upper end of the confidence interval.

If we abandon the truncation bias from the normal approximation in the parameter regime where the variance is small, using

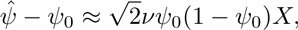

where *X* has standard normal distribution. Conditioned on*ψ̂* > 0 we have the conditioning bias

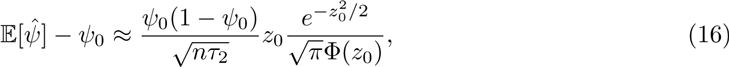

where 
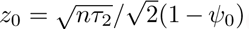
. If we truncate the estimates —— raising all negative estimates to 0 —— we get the truncation bias

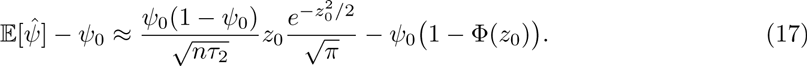

The argument for rejecting negative estimates appears reasonable, so long as the focus is only on the random-effects probability model that provided the original motivation. The parameter *ψ* is first introduced as a ratio of two variances, hence as a positive number. The denominator represents total variance and the numerator represents one component of variance, implying a ratio in [0, 1] if the two components are independent. (The question of whether *ψ* > 1 makes sense is a separate one, that we will not consider here.)

The common practice of restricting the maximum likelihood calculation to non-negative values, however, introduces bias that is well-known and may be serious for samples of moderate size, both when estimates are being truncated at zero or when negatives are being ignored. It is thus worth looking beyond original motivation to the actual structure of the GCTA model and considering what meaning negative parameter values might turn out to have.

The “variance attributable to additive genetic effects”, while an essential component of the genetic framework that motivates GCTA, has no place of its own in the statistical model itself. It appears nowhere in the observable data. As a statistical model of the data, the GCTA random-effects model is simply a subclass embedded in the multivariate normal model defined by the family of covariance matrices (2). The subclass need not be defined by *ψ* ≥ 0, rather than by the formally more natural constraint 
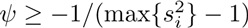
.

As described in section 4.1, the parameter for heritability here has the character of a correlation: the correlation between genotype similarity and phenotype similarity. The model invites us to select an estimate of *ψ* that will best match the genetic covariance between individuals to the similarity in their traits. High heritability means that individuals with similar genotype are likely to have similar trait values. Zero heritability means that genotypes tell us nothing about similarities in trait values. Negative heritability, then, could be perfectly sensible as a description of the data: It means that individuals with similar genotypes are likely to have more divergent trait values than those with highly disparate genotypes.

It is clear, then, that GCTA’s random-effects model of genetic causation can produce data that are best described by a negative *ψ*, hence by negative heritability. If we knew that the true parameter must be nonnegative, it would be hard to justify a negative estimate, rather than the estimate 0, which would have to be closer to the truth. Could a negative estimate ever be correct? Can we define a biologically plausible setting, of the same sort as the mixed-effects model that underpins GCTA, that naturally produces the data distribution defined by the full range of *ψ*, positive and negative, in the multivariate normal distribution?

In [14] J. B. S. Haldane described how the maternal-effect trait of neonatal jaundice could be said to display negative heritability: Because the disease results from maternal antibodies against a fetal antigen, it will not arise in a fetus whose mother herself experienced neonatal jaundice.^1^

Haldane calculates a negative heritability from a model that is specialized to the peculiar structure of neonatal jaundice. We show now that a simple model of gene-environment interaction naturally produces such a process under more general circumstances. We emphasize here that this does not involve any deviation from the multivariate normal distribution as defined; in particular, the effects of different SNPs remain independent, and there is no dependence between genotypes and “noise”.

Suppose we have a sequence of loci 1, …, *p* with linear reaction norms with regard to an environmental variable *ω*, which is assumed to have mean 0. We assume that environment and genes are independent. (A similar effect could be produced by gene-environment correlation, but we confine our consideration to gene-environment interaction.) For simplicity we assume that each locus is associated with two variants, denoted *Z_ij_* = 0 or 1 — perhaps because of dominance or other nonlinearity of interaction — so that the phenotype of individual *i* is

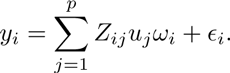

We define 
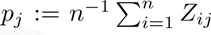
, the fraction of individuals with type 1 at site *j*, and assume that there is no linkage.

Consider two randomly chosen individuals, *i* and *i^′^*, whose genotypes agree in *k* sites and disagree in *p* − *k* sites. Conditioned on **u** and *Z*, we have

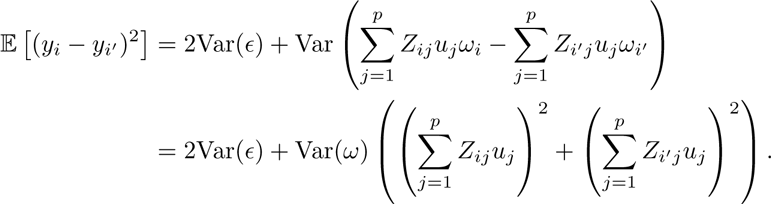

Now suppose we average over all the ways of agreeing in *k* sites. Conditioned on *j* being a site of agreement, the probability of both having type 1 is 
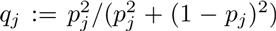
. Conditioned on *j* being a site of disagreement, 
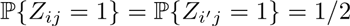
. Thus

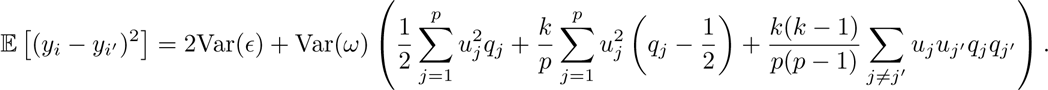

If the *q_j_* are significantly larger than 1/2 — meaning that the effective alleles are more common than the neutral alleles — and if the *u_j_* all have the same sign — meaning that the different alleles that affect *y* tend to push in the same direction in the same environment — then this will be strictly increasing in *k*, meaning that individuals with more similar genotypes will tend to have more disparate trait values. Even if the *u_j_* are only i.i.d. samples from some distribution, there is at least a 1/2 chance that the last term will be positive, so there would still be strictly negative heritability.

## 5. Model misspecification

### 5.1. Basic principles

Models that function well when fitted to data sampled from the correct distribution may produce unpredictable and unintuitive results when applied to data generated by a different albeit analogous distribution. Much of the dispute between [20] and [55] centers around the question of whether the specification of GCTA allows for linkage disequilibrium, and whether population stratification is adequately accounted for. [20] also considers, by means of subsampling real data, the question of whether the choice of SNPs to be investigated — as a subset of the full complement of SNPs — increases the variance of heritability estimates in ways that the standard analysis fails to capture.

The question that needs to be asked is this: Given the simplifications that we know underly the GCTA model, should we expect approximately sensible inferences to follow from approximately well specified data? We consider three types of deviation from the model:

- Shared environment;
- Measurement error;
- A small number of causal loci that are responsible for the influence of genes on phenotype, while the large majority of SNPs have no influence.

### 5.2. Shared environment

This problem is well known, and a focus of significant attention. It has long been recognized that large and small singular values of the genotype matrix are associated with potential confounding of genetic and environmental influences. Large singular values arise from stratified populations, reflecting geographical or ethnic differences that may be associated with trait differences not directly caused by the genetic differences themselves. Small singular values tend to arise from small clusters of related individuals, who are likely to be correlated in their environments as well. (In the extreme case, *C* clones produce *C* − 1 zero singular values and one singular value of size *C*.) The usual practice of working with the GCTA model recognizes these problems: from the outset, population stratification has been addressed with principal component analyses and cryptic relatedness by removing samples with suspiciously high kinship [53].

We have ignored environmental effects here (except in the discussion of negative heritability in section 4.3), focusing on situations where all assumptions of the GCTA model hold exactly. The one thing we have to add to this discussion is to point out that the behavior of models such as GCTA depends entirely on the distribution of singular values of the genotype matrix, and thus any confounding must manifest itself through these singular values. In particular, any excess variance among the singular values relative to the known limiting form in the independent setting — which models i.i.d. genotypes that, by construction, cannot be confounded — must come from latent structure between samples or between loci. Inter-sample structure almost inevitably allows the intrusion of shared environment. That is, related samples present a tradeoff: they increase spectral spread, decreasing the heritability estimator’s variance, but expose GCTA to potential confounding bias. Therefore, if one is convinced that the latent inter-sample structure is benign — possibly, for example, in laboratory animals or carefully controlled twin studies — the additional spectral variance improves heritability estimates.

This also explains apparent peculiarities in distribution of squared singular values in the independent setting described in Section 6. First, it has disappointingly small variance because there is no latent sample structure. More importantly, the problem is exacerbated when *p* grows larger, as relatedness that emerges due to chance from a small number of i.i.d. draws will converge asymptotically to zero, the expected relatedness in i.i.d. data. This is something like genome-wide Mendelian randomization, and one expects these purely exogenous genotype effects to cancel out as the number of independent genotype contributions increases.

### 5.3. Measurement error

We note here that simple measurement errors do not cause any unusual problems for the mixed-effects model; as one would expect, it simply biases the estimates of heritability downward. We look here at two types of error: Independent additive error in measuring phenotypes, and independent misidentification of SNPs. Adding an independent measurement error to the phenotype simply increases the variance of the noise term *ϵ*, so it is equivalent to the same model with a lower value of *ϕ* (or *ψ*).

Misidentification of SNPs is slightly more complicated. Suppose that instead of observing *Z*, we observe *Z̆* = *Z* + *Z̃*. We assume the entries of *Z̃* to be independent of each other, and of the noise, with expectation 0. (They obviously can’t be independent of the entries of *Z*, but *Z* is taken to be fixed, not random.) They are nonzero with probability *π*, which we assume is close to 0. Then the phenotypes will satisfy

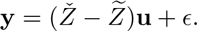

Applying the singular value decomposition *Z̆* = *U*diag(*s_i_*)*V*\*, we get

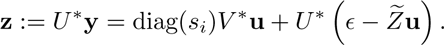

The term *ϵ* − *Z̃***u** is approximately a vector of independent normal random variables, with mean zero and variance 
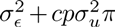
 where *c* depends on the distribution of e *Z̃*. Its covariance with diag(*s_i_*)*V\****u** will be on the order of *V*Z̃*, which has expectation 0 (averaged over realizations of *Z̃*), and should typically be on the order of 
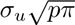
, meaning that the correlations are small as long as *pπ* is large.

We may conclude, then, that the model with occasional misidentification of SNPs is very much like the model with increased noise in the phenotype measurement, with a downward bias in heritability estimates proportional to the error probability.

### 5.4. Causal loci

The usual practice of working with the GCTA model recognizes that the genetic effect saturates as the number of SNPs sampled increases [27, 30, 28, 60, 11, 10]. This is generally attributed to the increasing amount of linkage to the (possibly unobserved) causal loci. Here and in the following section we analyze the effect of applying GCTA in a situation where there is a small number of causal SNPs, either a small subset of the observed SNPs (section 5.4.1) or an unobserved set that may be linked to the observed SNPs (section 5.4.2).

#### 5.4.1. Observed causal SNPs

Suppose that the genetic effect on **y** is produced by a small number *k* ≪ *n* of SNPs. Other SNPs will be linked to these, thus being indirectly correlated with **y**. We may represent this as a slightly modified version of the standard GCTA model by assuming that there is a subset *η* ⊂ {1, …, *p*} of causal SNPs, and that these causal SNPs have i.i.d. normal effects, with mean 0, conditioned on the sum of their squares being a fixed number 
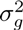
. We will also think of *η* as a *p*-dimensional vector with 1 in the positions corresponding to the causal SNPs and 0 elsewhere.

We write the noise variance as

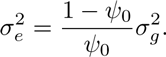

(The “true heritability” is naturally identified with *θ*‖**u**‖^2^; when all but a small number of the components of *u* are zero, this will not necessarily be very close to *ψ*_0_ unless we impose this as a condition.) When we think of the set of causal SNPs *η* as being fixed we will call this the *causal-SNP GCTA* model (or CS); when we think of *η* as being a uniform randomly selected subset we call it the *random causal-SNP GCTA* model (or RCS).

Conditioned on a fixed *η*,

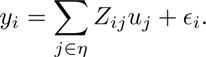

The MLE will converge, as described in [49], to the closest fit (in the Kullback–Leibler sense) to the generating model. The question then is, what is the *ψ_*_* that maximizes the expected log likelihood? Equivalently, we seek the *ψ_*_* that solves

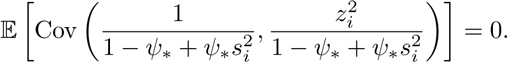

By linearity of covariances, this becomes

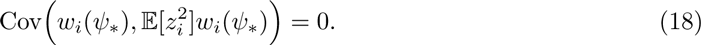

In the CS model (so, holding *η* fixed)

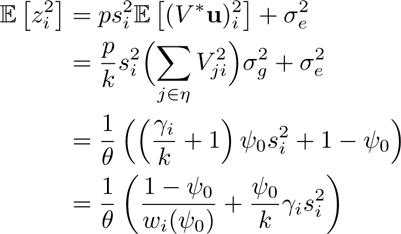

where

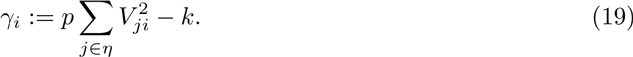

represents the deviation from expectation of the size of the projection of *η* onto the *i*-th right singular vector. We note for later that *γ_i_* has expectation zero (with respect to the choice of a random *η*), and is identically zero when *k* = *p*. We note as well that when *k* ≪ *p*, we will typically expect *γ_i_* + *k* to be distributed approximately like a chi-squared variable with *k* degrees of freedom.

The equation (18) for the value*ψ*, to which the estimate*ψ̂* will tend asymptotically, then becomes

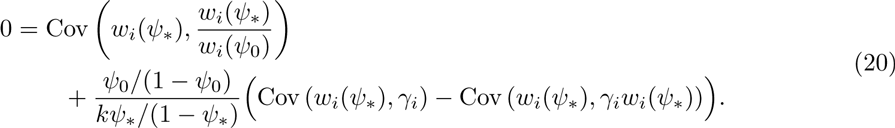

There are two ways we might use this equation. For a given choice of singular-value distribution and of true parameter *ψ*_0_, this equation defines *ψ_*_* as a function of (*γ_i_*). For a given genotype matrix we could compute the distribution of the *γ_i_* jointly with the phenotypes for a random choice of possible causal sites. In this way we could more efficiently simulate the effect of restricted causality on the heritability estimates.

Alternatively, we could use the assumption that the *γ_i_* are generically i.i.d. samples from a chi-square distribution. The expression on the right-hand side of (20) is a linear combination of these, hence is approximately normal. In fact, since we assume *n* is large, and the *w_i_*(*ψ_*_*) are bounded, the approximation should be extremely good. The variance of the sum is the sum of the squares of the coefficients, multiplied by the common variance 2*k*. Using the equality

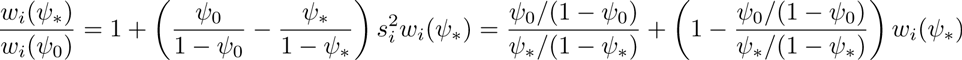

this yields

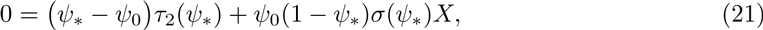

where *X*, defined as a function of the *γ_i_*, has standard normal distribution, and

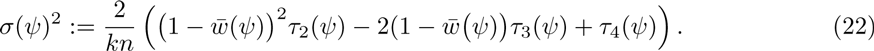

(Note that we have multiplied by *ψ_*_* — in fact, we have done so twice — so we are only considering positive solutions of the equation.)

We may now perform a perturbation analysis, on the assumption that the discrepancy between *ψ_*_* and *ψ*_0_ is small. If this is true, then we can obtain a first-order approximation for *ϵ*:= (*ψ_*_* − *ψ*_0_)/(*ψ*_0_(1 − *ψ*_0_)) by solving

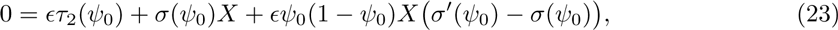

leading to

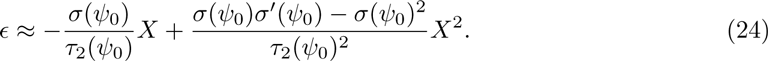

Assuming *σ*/*τ*_2_ and *σ′*/*τ*_2_ are both small, and that *τ*_3_ and *τ*_4_ are much smaller than *τ*_2_, we have the approximations

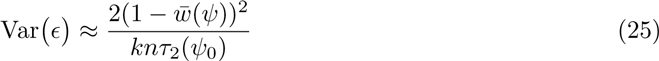

and

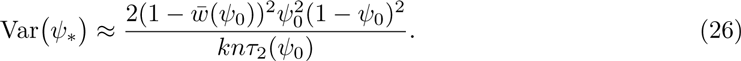

If we restrict attention to the independent setting, and assume that *μ* = *n*/*p* is very small, then we may draw on the results in Section 6 and solve the equation explicitly. We have *w̄* ≈ 1 − *ψ_*_* and 
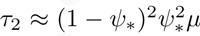
, while *τ*_3_ and *τ*_4_ are moderate multiples of *μ*^2^. We conclude that, to first order in *n*^−1^,

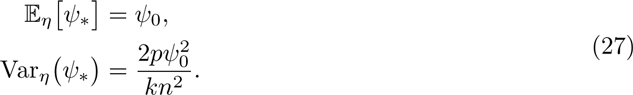

Regardless of their exact distribution, if the eigenvalues are sufficiently concentrated around 1 that *τ_k_* ≪ *τ*_2_, then we will have a ratio of variance due to random selection of causal SNPs to the variance due to random genetic effects and genetic phenotypes (the variance considered in standard analyses, given in (14)) of

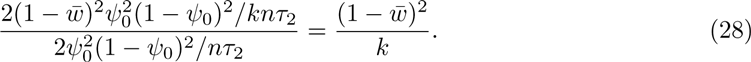

Thus, it may be assumed to be negligible — relative to the uncertainty generally acknowledged in the standard analysis — as long as the phenotype is influenced by several tens of SNPs, but to increase the uncertainty substantially when fewer than 10 SNPs are involved. This effect will be exacerbated when *w̄* is small, which will be the case when the heritability is high.

To illustrate this effect, we conducted simulations in the independent setting. We took *p* = 5000 SNPs, and defined them to have minor allele frequencies independently chosen, uniform on [0.05, 0.5]. We then simulated genotypes for *n* =1000 or 2000 individuals by independently assigning a random number of minor alleles to each individual and site, according to the binomial distribution with the appropriate MAF. The genotypes corresponding to each site were then normalized to have mean 0 and variance 1. This was our genotype matrix *Z*.

We then selected a random subset of sites to be the causal ones, and simulated 1000 phenotypes for each individual, with heritability either 0.5 or 0.8. The heritability was then estimated, according to the standard MLE procedure described in section 3. We then repeated this procedure for 100 different random choices of the causal SNPs.

The results are summarized in Figure 3, which shows the variance of the genotype-mean simulation outcomes (points), and compares it to the theoretical prediction of (26) (continuous curves) as *k* varies. Overall, the theoretical curve is fit very well, though the fit is worse when the variance between different genotypes is small (*k* large), making it hard to separate from the much larger phenotype variance.

**Fig 3.**
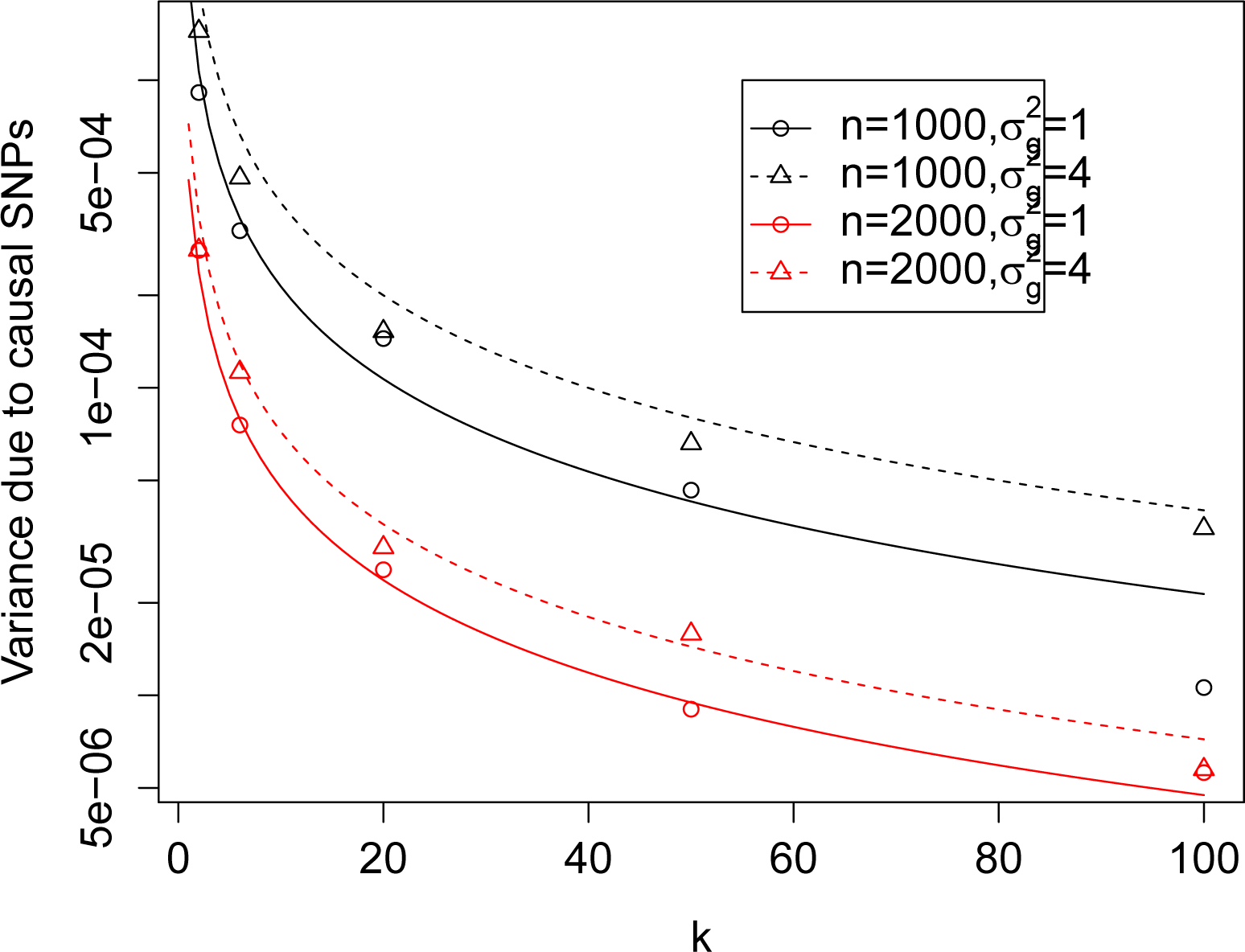
Estimated variance of average heritability estimate for 1000 random phenotypes, from each of 100 randomly selected subsets of k causal SNPs.

#### 5.4.2. Unobserved causal SNPs

It is generally assumed that the SNPs that directly influence the phenotype *y_i_* are not actually among the *p* SNPs that have been measured, the problem of “untagged variation”. Clearly we cannot assume that the estimate of *ψ* will be unbiased. In the extreme case where the causal SNPs are independent of the observed SNPs, of course the expected estimate of *ψ* will be 0. In general we expect to see a downward bias, since the residual uncertainty about the causal SNPs will act like regression measurement error, deflating our estimate of the regression slope, which is heritability. While this has been remarked qualitatively both for GWAS [34] and GCTA [53, 23] heritability estimates, we are not aware of a formal derivation of the effect of untagged variation on heritability estimates.

Intuitively, it makes sense that the estimation will be as good as the best possible imputation of the causal SNPs. Of course, if we knew which were the causal SNPs we could simply include them in the sample — either the measured, or the imputed values. We are assuming, though, that there is no information about the causal SNPs, which are not in the panel.

More to the point, there is no inherent meaning to “best imputation” outside the context of a particular probabilistic model generating the genotypes. For a given collection of observed and unobserved (but causal) genotype data there is an answer to the question, what is the bias in the heritability that would be estimated if we calculated the MLE from the observed genotypes? We write down this answer formally in (30), but do not see any meaningful interpretation of this formula. If we embed the genotypes in a probabilistic model, on the other hand, we are able to discuss the distribution of the unobserved bias understood as a random quantity, just as we did in section 5.4.1.

The model is exactly the same as in section 5.4.1, except that the *k* causal SNPs are not among the *p* observed SNPs, so the model now includes *p + k* SNPs in total. (We do not assume independence, so this model includes the observed-causal model as a special case, if we simply make the causal columns to be copies of some of the observed columns.)

We consider two probabilistic models:

1. The causal sites are a random sample of all sites, which are then not included in the observed genotype matrix.
2. The causal genotype matrix is generated by a linear relation

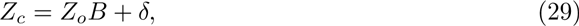

where *Z_o_* are the genotypes at the *p* observed loci (*Z* above) *B* is a fixed *p* × *k* matrix, and *δ* a random *n* × *k* matrix with mean-zero independent entries, such that the entries in column *ℓ* have variance 
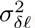
. We write

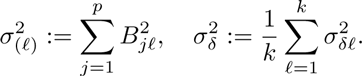

We assume that *B* is approximately sparse — all but a small fraction of entries are negligibly small — with no more than one non-small entry in any row. That is, there is a small number of observed sites that yield nearly all the information about an individual’s causal SNPs, and these linked sites are distinct for different causal SNPs. We also assume that

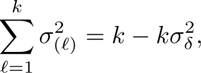

which is simply a matter of ensuring that *σ*^2^ is actually the additive genetic variance.

The first model will not be discussed in detail here. It is a subject for empirical exploration, only a slight formalization of the simulation tests commonly applied as in [53] to validate GCTA. The second is somewhat unsatisfactory, as it produces abstract “genotypes” that are unlike the real 0,1,2 genotypes produced in real experiments. We describe it here to illustrate how the behavior of such models may be rigorously analyzed, though a more realistic version would be technically more demanding.

Taking *U∑V*\* now to be the singular-value decomposition of 
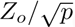
, we compute the rotated phenotypes

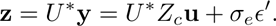

where *ϵ′*:= *U*ϵ* is another vector of i.i.d. standard normal random variables. Thus

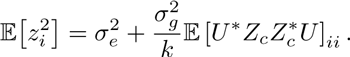

The estimator *ψ_*_* will satisfy

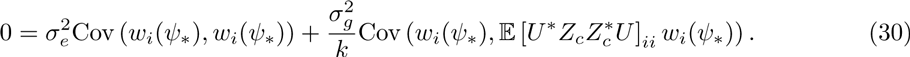

At the moment there is not much we can say further about the first model. We do not know of any general results relating the SVD of a matrix to the SVD of a random sample of its columns. It would be possible to investigate (30) through simulation.

We can say more about the second model. We have

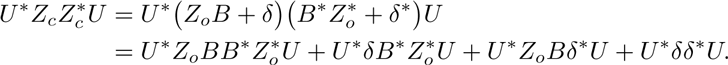

Since the rows of *δ* are uncorrelated with mean 0 we have 
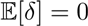
 and

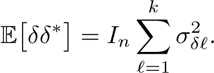

By definition, 
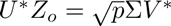
. Thus

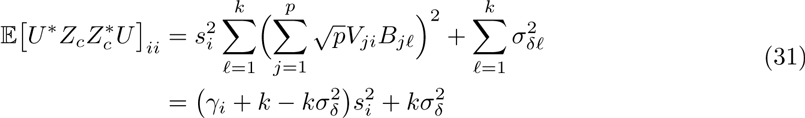

where

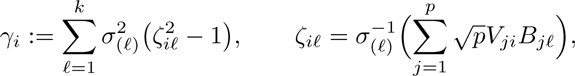

and so

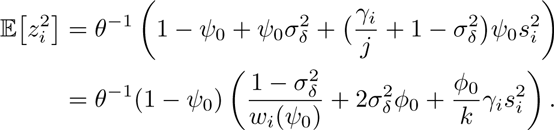

Thus (18) becomes

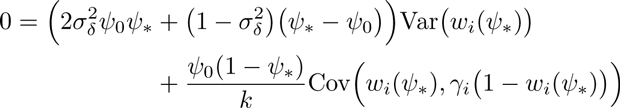

By the same sort of calculation as before, we may write

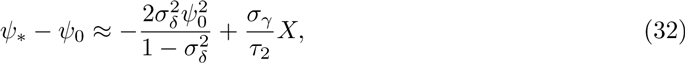

where *X* is approximately standard normal (as *B* varies over different permutations of possible causal SNPs) and

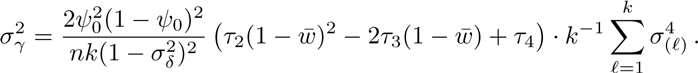

We may draw three conclusions:

1. When *σ_δ_* is not zero (or very small) — that is, when the causal SNPs are not completely determined by the observed SNPs — there is a negative bias in the heritability estimate, roughly proportional to 
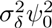
, which is a measure of untagged variation.
2. The correction for “imperfect LD” from [53] shrinks *A* toward the identity with *βA* + (1 − *β*)*I* or, equivalently, inflates the heritability estimate from *ψ̂* to 
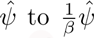
 [12]. *β* is estimated under biologically motivated “strong assumptions” and may not be necessary for whole genome sequencing or imputed data [52]. However, even under these assumptions, no value of *β* can generally correct the bias for all *ψ*. Specifically, even if we ignore the bias due to *X* — i.e. due to randomness in causal variants/*B* — the bias in (32) scales with *ψ*^2^, not *ψ*_0_.
3. When *σ_δ_* is zero this includes the situation of section 5.4.1, if *B* is a binary matrix with only ones and zeros, so that each causal SNP is a copy of an observed SNP. The formula (32) generalizes the calculation from the previous section, so that we see that the added uncertainty (or random bias) decreases when the information about each causal SNP is split up among multiple observed SNPs.

## 6. Singular Values

We now present formulas for moments relevant to estimator bias and variance for special cases of the distribution of squared singular values on which the GCTA estimates depend. We consider first the independent setting, followed by several stylized models for a stratified setting. Whereas our general treatment has only assumed a normalization of the total sum of squares of the elements of the genotype matrix, for these special cases we assume — as is usual in applications of GCTA — that each column of the genotype matrix has been normalized to have unit variance as well as zero mean. The methods of this section can be extended to cases with dispersion in column variances, as well as to genotype matrices with linkage disequilibrium, but we do not pursue these extensions here.

There is a closed-form limiting expression for the empirical measure of the singular values in our independent setting as *n* grows large for fixed *μ* = *n*/*p*. It was discovered by Marcenko and Pastur [35] and independently by Mallows and Wachter (see [33]). We use it in the generality established by Wachter [47].

The theorems provide for almost-sure convergence to a deterministic limit. Any fixed genotype matrix *Z* will have singular values with an empirical measure close (to order 1*/n*) to this limit, if *Z* falls within a set of matrices that would have probability one in an ensemble of random matrices with independent elements. Recall that the column variances of *Z* are normalized to be close to unity.

In order to find expressions for the empirical moments across *i* of *w_i_*(*ψ*) as functions of *ψ* and *μ* = *n*/*p* define the Stieltjes Transform for complex *ζ* away from the real interval [*a, b*] by

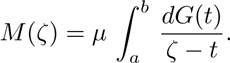

Here *t* = *s*^2^ stands for the eigenvalues corresponding to the singular values, and *dG* is the limiting empirical/measure of the eigenvalues, concentrated on the interval [*a*, *b*] where 
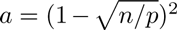
 and 
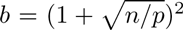
. Conversion of the formulas for singular values to eigenvalues requires removing mass 1 − 2*μ* conventionally placed at zero and rescaling by *μ* so that *dG* has unit mass on [*a*, *b*].

The average of *w_i_* is given by the value of *ζM*(*ζ*)/*μ* and the average of 
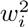
 by the value of −*ζ*^2^*M′*(*ζ*)/*μ* when we plug in *ζ* = −(1 − *ψ*)/*ψ* = −1/*ϕ*, making 1/(1 − *ζ*) equal the heritability *ψ*. Averages of higher powers of *w_i_* are given by expressions in higher-order derivatives of *M*(*ζ*)/*μ*. Equation 2.1.1 of [47] shows that *M*(*ζ*) is the solution vanishing at infinity to the quadratic equation

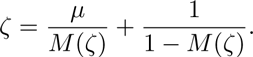

The solution can be written

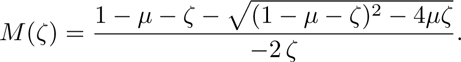

Here the sign on the square root is chosen to agree with the sign on 1 − *μ* − ζ in order to make *M*(*ζ*) vanish at infinity and to make *ζM*(*ζ*) approach *μ* at infinity.

The expression for *M* (*ζ*)/*μ* containing the square root can be differentiated in closed form, and exact expressions for the moments of *w_i_* follow from substituting 1/(1 − *ζ*) = *ψ* and −*ζ* = (1 − *ψ*)/*ψ*. For practical purposes it is helpful to expand *M*(*ζ*)/*μ* in powers of *μ*. The coefficients conveniently arrange themselves in powers of (1−*ζ*)^−1^, streamlining differentiation and calculation of uncentered moments of any order for *w_i_*.

Specifically, for small *μ* and *ζ* either off the real axis or outside [*a, b*], we have

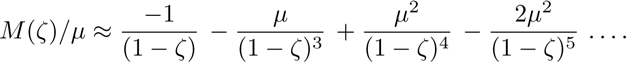

When 1/(1 − *ζ*) = *ψ*, and −*ζ* = (1 − *ψ*)/*ψ*, evaluating *ζM*/*μ* we find that the mean over *i* of *w_i_* is given up to second order in *μ* by

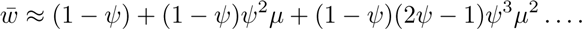

Differentiating *M* by *ζ* yields

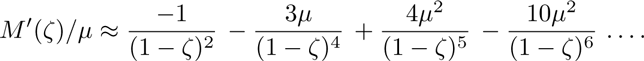

It follows that the variance over *i* of *w_i_* is given by

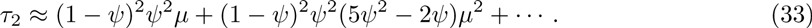

We also have

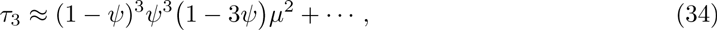

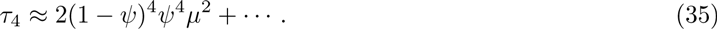

Higher-order moments follow by successive differentiation.

Central moments of the eigenvalues themselves around their mean of unity can be found by collecting terms in powers of 1/(1 − *ζ*). The variance is *μ*, and to second order in *μ* the third central moment is +*μ*^2^ and the fourth central moment 2*μ*^2^. Evaluating *M*(*ζ*)/*μ* at *ζ* = 0 shows that the mean of the reciprocal eigenvalues is 1/(1 − *μ*), and successive differentiation reveals a variance of *μ*/(1 − *μ*)^3^ and a third central moment of 2*μ*^2^/(1 − *μ*)^5^ for the reciprocal eigenvalues.

For the stratified setting, a great variety of scenarios come under consideration. Stratified populations are expected to have genotype matrices with some or many eigenvalues substantially larger than those in the independent setting, and large eigenvalues entail large numbers of small ones, since the mean is unity.

Among the simplest stylized models for eigenvalues arising in a stratified population, one posits eigenvalues concentrated at two reciprocal values 1/*β* and *β* with weights *β*/(*β* + 1) and 1/(*β* + 1). This two point-mass distribution, with its mean of unity, has variance (*β* − 1)^2^/(*β* + 1), close to *β* when *β* is large. Moments over *i* of *w_i_* come out to be

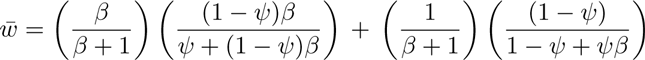

and

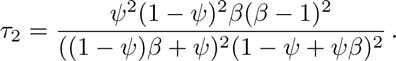

For large *β*, the mean of *w_i_* is close to 1 and the variance of *w_i_* is close to 1*/β*. In contrast to the independent setting, where the variance of *w_i_* is roughly proportional to the eigenvalue variance, in this stylized model for the stratified setting, the variance of *w_i_* is roughly inversely proportional to the eigenvalue variance. Implications for bias and variance for GCTA estimates have been described in section 3.3.

## 7. Discussion

In the simple setting when assumptions are satisfied, we have shown that threats to the accuracy of GCTA estimates arise not from bias but from potentially large standard errors. Our findings run counter to some recent criticisms of GCTA.

We have also evaluated two sources of bias in more complicated settings – bias from exclusion of negative estimates of heritability and bias from fixed but unknown structured subsets for causal SNPs. Both can be substantial in principle. But neither seems likely to be disabling in practice. As for negative estimates, we have argued that they remain meaningful within the statistical specification for GCTA; retaining them avoids that bias. As for subsets of causal SNPs, we have argued that kinds of structure amplifying bias are likely to be the exception rather than the rule.

Standard errors for GCTA estimates of heritabilities depend on the dispersion in the squared singular values of the genotype matrix. In this regard, an idealized baseline case, the independent setting, is something like a worst-case scenario. Here, drawing on eigenvalue theory, we have given explicit expansions for standard error, as well as bias, in terms of the ratio of respondents to SNPs. Implications of our formulas have been reviewed in Section 3.4. Departures from the independent setting which augment the dispersion of the squared singular values can improve the statistical properties of the estimates, but only up to a point and only at a cost in substantive interpretability. Linkage disequilibrium augments dispersion. Unexpunged population stratification augments dispersion. Their relative roles and their signatures are not yet clear. A priority for future research is analysis of empirical genotype matrices and their sets of singular values, the natural next step in elucidating the statistical properties of random effects models for genetic heritability.

## Footnotes

1 We thank Jonathan Marchini for pointing out this reference to us.

